# Type I interferons govern Zika virus resistance of human macrophages and microglia

**DOI:** 10.1101/2022.03.09.483671

**Authors:** Aidan T. Hanrath, Catherine F. Hatton, Florian Gothe, Cathy Browne, Jane Vowles, Peter Leary, Simon J. Cockell, Sally A. Cowley, William S. James, Sophie Hambleton, Christopher J. A. Duncan

## Abstract

The role of the human type I interferon (IFN-I) system in restricting Zika virus (ZIKV) is uncertain. Here, genetic and pharmacological ablation of IFN-I signalling enhanced ZIKV replication and cytopathicity in macrophages and microglia, key cells in ZIKV transmission and pathogenesis. Thus, despite the extensive IFN-I countermeasures employed by ZIKV, IFN-I dictates the outcome of infection in macrophages. Therapeutic manipulation of the IFN-I system may bring clinical benefit in ZIKV.

## Main text

Type I interferons (IFN-I) are essential to antiviral immunity^1,2^. Signalling through a ubiquitously expressed receptor formed of *IFNAR1* and *IFNAR2*, IFN-I activates a JAK-STAT signalling pathway culminating in the expression of hundreds of interferon-stimulated genes (ISGs) that mediate its antiviral properties^2^. Zika virus (ZIKV) is a neurotropic flavivirus of global public health importance. ZIKV epidemics are associated with pregnancy loss and neurological syndromes including microcephaly, meningoencephalitis and Guillain-Barré syndrome ^3^. ZIKV, like other human viral pathogens, has evolved multiple strategies to subvert IFN-I restriction in human cells^4,5^. Failure to achieve this in other species limits ZIKV host range^6^, mandating studies in human systems. Understanding the molecular interactions between ZIKV and the human IFN system is critical to understand pathogenesis and guide therapeutic strategies^5^. Studies have yielded inconsistent findings concerning whether ZIKV induces an endogenous IFN-I response in human cells^4,5,7-15^, and none have directly assessed its functional relevance in ZIKV restriction.

We addressed this question in macrophages, a key target cell attributed to both protective and pathologic roles in ZIKV^16^. We employed a validated model of human macrophages derived from induced pluripotent stem cells (iPS-Mϕ)^17^, generated from an IFNAR2-deficient^1^ patient (*IFNAR2*^PT^ Fig. 1a, Fig. S1a-c) alongside two healthy controls (*IFNAR2*^WT^). *IFNAR2*^PT^ iPS-Mϕ exhibited a profound defect of IFN-I signalling (Fig. S1d). Flow cytometry analysis confirmed appropriate expression of macrophage surface markers and phagocytic function (Fig. S1e-f). Both *IFNAR2*^PT^ and *IFNAR2*^WT^ iPS-Mϕ produced IFNs upon infection with Asian lineage ZIKV H/FP/2013 (ZIKV^FP^), assessed by RT-PCR analysis at 24 hours post-infection (h.p.i.) (Fig. 1b). Immunoblot analysis (48h.p.i.) demonstrated expression of phosphorylated STAT1 accompanied by the ISGs RSAD2 and ISG15 in *IFNAR2*^WT^, but not *IFNAR2*^PT^ iPS-Mϕ, reflecting the defect of IFN-I signalling in the latter (Fig. 1c).

**Figure 1.**
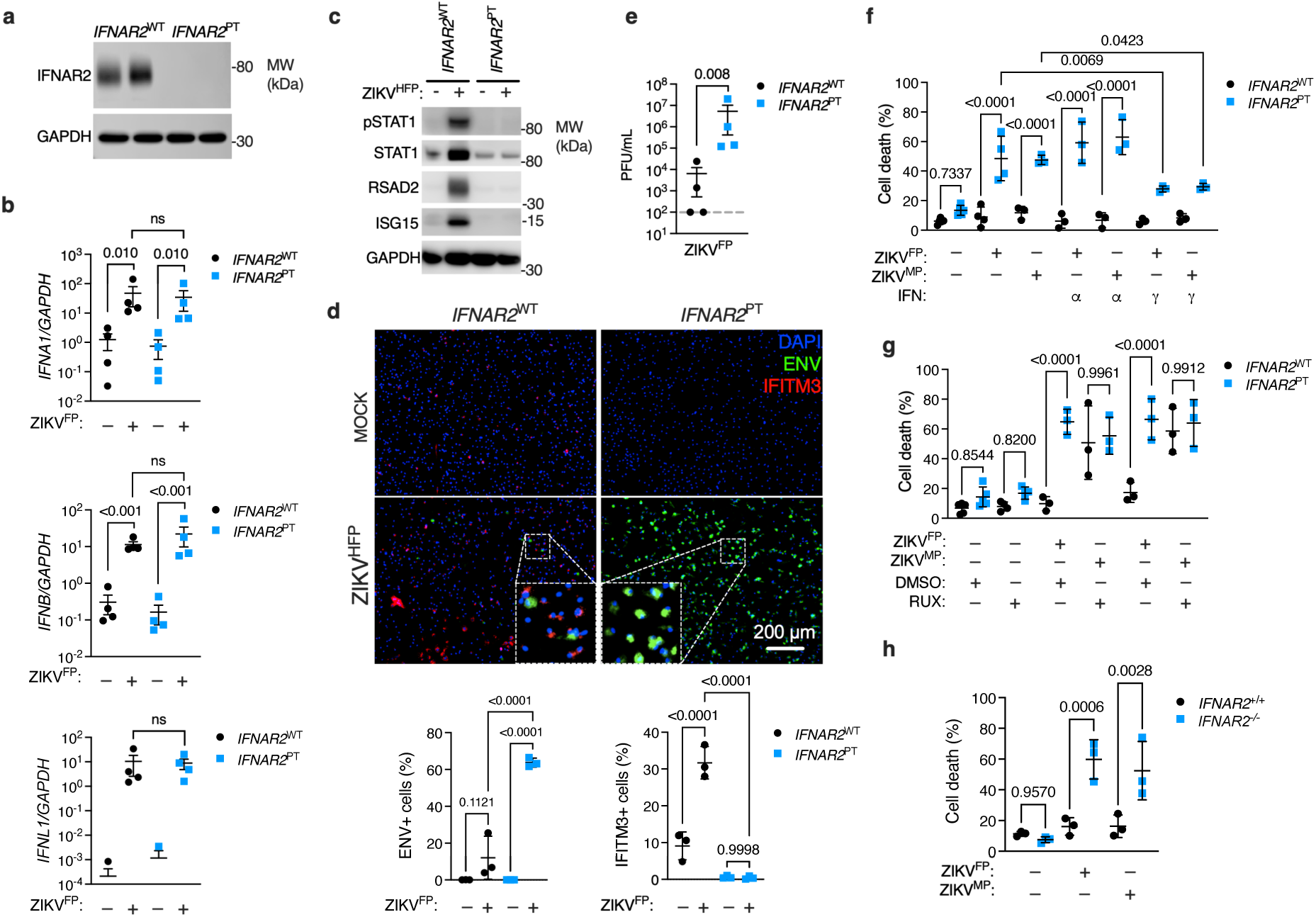
Enhanced ZIKV replication and cytopathicity in IFNAR2-deficient macrophages. (a) Immunoblot of IFNAR2 and GAPDH in *IFNAR2*^WT^ (WT2) and *IFNAR2*^PT^ (clone 11) iPS-Mϕ, representative of n=3 repeats. (b) RT-PCR quantification of *IFNB, IFNA1* and *IFNL1* relative to *GAPDH* (24h.p.i. ZIKV^FP^ MOI=10.0, n=4 repeat experiments in *IFNAR2*^PT^ [clone 6] and *IFNAR2*^WT^ [WT2]). (c) Immunoblot of pSTAT1, STAT1, RSAD2, GAPDH and ISG15 in *IFNAR2*^WT^ (WT2) and *IFNAR2*^PT^(clone 11) iPS-Mϕ (48h.p.i. ZIKV^FP^ MOI=1.0), representative of n=3 repeats. (d) Immunofluorescence analysis of ZIKV ENV and IFITM3 expression (48h.p.i. ZIKV^FP^ MOI=1.0, representative images and quantification of n=3 repeats). (e) Plaque assay of ZIKV HFP (48h.p.i. ZIKV^FP^ MOI=0.001, n=4 repeats). (f) Cell viability assay (72h.p.i. ZIKV^FP^ or ZIKV^MP^ MOI=1.0, n=3-4 repeats). (g) As in (f), in ruxolitinib (10 μM) or vehicle (DMSO)-treated *IFNAR2*^WT^ iPS-Mϕ (WT1 and WT2, n=3-4 repeats). (h) As in (f), in *IFNAR2*^-/-^ (G2, B5_2) and isogenic *IFNAR2*^+/+^ (G8, F8) iPS-Mϕ. Unless stated, all data are mean±SD of n=3 independent repeats in *IFNAR2*^PT^ (clones 6 and 11) and *IFNAR2*^WT^ (WT1 and WT2), with comparisons by t test (d-e) or ANOVA with Sidak’s test (f-h).

This was associated with a substantial enhancement of ZIKV replication in *IFNAR2*^PT^ iPS-Mϕ, quantified by various methods, including analysis of viral Envelope (ENV) expression and plaque assay of infectious particle release (Fig. 1d-e, Fig. S2). This phenotype was independent of viral origin, as it also followed African lineage MP1751 infection (ZIKV^MP^ Fig. S2). Expression of the ISG IFITM3 was significantly elevated in *IFNAR2*^WT^, but not *IFNAR2*^PT^ iPS-Mϕ, and was seen predominantly in uninfected bystander cells, consistent with paracrine IFN-I signalling. Accompanying heightened ZIKV replication in IFN-I incompetent cells, we observed morphological abnormalities suggestive of cell death, with progressive membrane blebbing and cell shrinkage by 72h.p.i (Fig. S3a) accompanied by immunodetection of cleaved caspase 3 (Fig. S3b), indicative of apoptosis^14^. These data indicated that failure of IFN-I mediated control led to cytopathic effects (CPE) in *IFNAR2*^PT^ iPS-Mϕ, contrasting with the resistance of WT macrophages to ZIKV CPE^7,10,11^. Quantifying this in a live cell viability assay, we confirmed that ZIKV CPE was significantly enhanced in *IFNAR2*^PT^ but not *IFNAR2*^WT^ iPS-Mϕ (Fig. 1f). Treatment of *IFNAR2*^PT^ iPS-Mϕ with IFN*γ;* partially rescued this phenotype.

To verify whether this vulnerability could be recapitulated by IFNAR blockade in WT cells, we adopted two complementary approaches: first, we used the pharmacological JAK inhibitor ruxolitinib; second, we used CRISPR/Cas9 to delete *IFNAR2* in both control lines, selecting single cell clones and generating *IFNAR2*^-/-^ and isogenic *IFNAR2*^+/+^ iPS-Mϕ (Fig. S4). These approaches recapitulated CPE and enhanced ZIKV infection seen previously in *IFNAR2*^PT^ iPS-Mϕ (Fig. 1g-h, Fig. S4g), confirming their association with IFN-I incompetence.

To explore the global transcriptional response to ZIKV, we undertook RNA-sequencing of *IFNAR2*^-/-^ and isogenic *IFNAR2*^+/+^ iPS-Mϕ clones at 24h.p.i. Principal component and pathway analyses revealed the dominant contribution of IFN-I signalling to the ZIKV response (Fig. 2a-b). Of 492 genes significantly differentially expressed (DE) by *IFNAR2*^-/-^ and *IFNAR2*^+/+^ iPS-Mϕ upon ZIKV infection, 467 (95%) were ISGs (http://interferome.its.monash.edu.au/interferome). Visualisation of significant context-specific macrophage ISGs^18^ highlighted this contrast (Fig. 2c). ZIKV reads were also significantly enriched in *IFNAR2*^-/-^ iPS-Mϕ (Fig. 2d), consistent with previous findings. Finally, we investigated the relevance of IFN-I to antiviral defence of microglia: brain-resident macrophages implicated in ZIKV neuropathology^11,19^. Employing a validated method for differentiating microglia-like cells (iPS-MGL)^20^ (Fig. S5), we observed the same behaviour in IFNAR2-deficient iPS-MGL, with heightened ZIKV replication and CPE (Fig. 2e-f).

**Figure 2.**
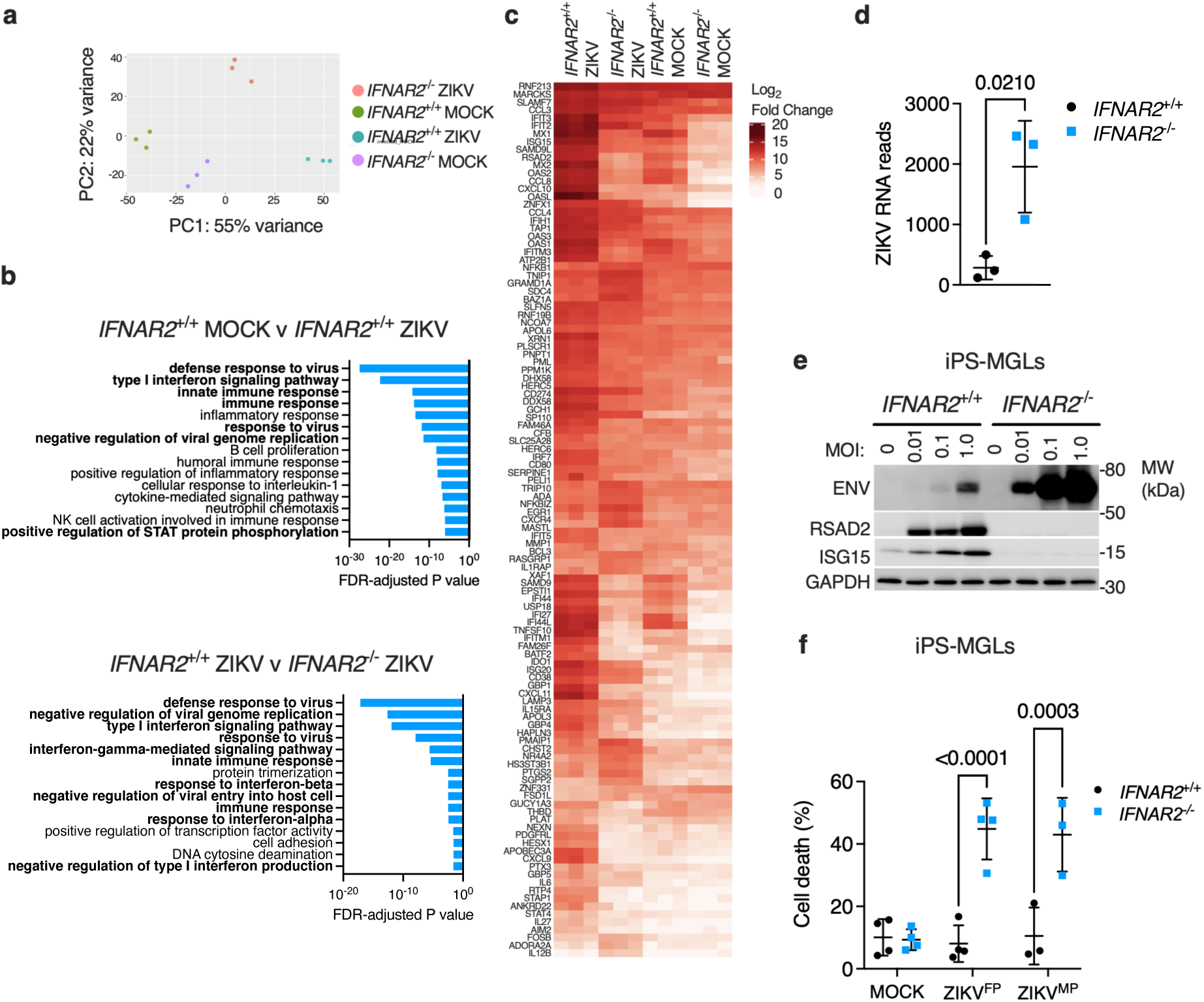
IFN-I signalling dominates the transcriptional response to ZIKV. (a) Principal component analysis of RNA-seq data (24h.p.i. ZIKV^FP^ MOI=1.0, n=3 independent repeats in isogenic *IFNAR2*^-/-^ [B5_2] and *IFNAR2*^+/+^ [F8] iPS-Mϕ). (b) Gene ontology analysis (FDR<5%) in mock v infected *IFNAR2*^+/+^ iPS-Mϕ (top) and *IFNAR2*^-/-^v *IFNAR2*^+/+^ iPS-Mϕ (bottom). Relevant pathways in bold. (c) Macrophage-specific ISGs (Log_2_FCϕ3, FDR<5%, colour intensity reflects Log_2_ FC). (d) ZIKV reads (mean ±SD, t test). (e) Immunoblot of ENV, RSAD2, ISG15, GAPDH in ZIKV^FP^-infected *IFNAR2*^+/+^ (F8) and *IFNAR2*^-/-^ (B5_2) iPS-microglia-like cells (iPS-MGLs 72h.p.i., representative of n=3 repeats in *IFNAR2*^PT^ [clones 6 and 11] and *IFNAR2*^WT^ [WT2]). (f) Cell viability assay (72h.p.i. ZIKV^FP^ or ZIKV^MP^ MOI=1.0, mean ±SD, n=3-4 repeats, ANOVA with Sidak’s test) in *IFNAR2*^-/-^ (B5_2), *IFNAR2*^PT^ (clone 6), *IFNAR2*^+/+^ (F8) and *IFNAR2*^PT^ (WT2) iPS-MGLs).

There were hints in the literature that human IFN-I participates in ZIKV resistance. For instance, ablation of AXL attenuated ZIKV replication in astrocytes by enhancing IFN-I signalling^8^. Similarly, deletion of *ISG15*, another negative regulator of IFNAR signalling, reduced ZIKV replication in iPS-derived neural progenitors^13^. Our data substantially extend this indirect evidence, showing that IFN-I is decisive in macrophage resistance to ZIKV, despite extensive viral countermeasures^5^.

Macrophages play an essential role in vertebrate host defence against microbes^16^. Many successful human pathogens, including HIV, have evolved sophisticated mechanisms to subvert macrophages - exploiting this cellular niche for survival and/or transmission^14^. ZIKV is thought to seed sanctuary sites such as the brain, testes or placenta via macrophages^7,10,11,19^, exploiting their resistance to CPE. Our data reveal that this resistance owes to robust IFN-I-mediated immunity, casting new light on the role played by these cells in ZIKV defence. The idea that macrophages might act in a protective capacity, as antiviral sentinels and/or effectors, is supported by recent findings in animal models^21,22^.

## Data availability

Source data are provided with this paper. Source data includes uncropped blots, all raw quantitative data and PFU counts. The results of differential expression analysis of RNA-seq data are included as a supplementary dataset S1. RNA seq data are available GEO (Accession: TBC).

## Acknowledgements

ATH was funded by an NIHR Academic Clinical Fellowship (ACF-2018-01-004) and the British Medical Association Foundation. FG is supported by the Munich Clinician Scientist Program at LMU (FoeFoLe^plus^) and received fellowships from the Bubble Foundation as well as the Care-for-Rare Foundation. CFH is funded by an MRC studentship [MR/N013840/1]. SH and CJAD are funded by the Wellcome Trust [207556/Z/17/Z and 211153/Z/18/Z respectively]. iPS cell derivation and gene editing was carried out at the James Martin Stem Cell Facility, which receives financial support from the Oxford Martin School (LC0910-004). We are grateful to Jon Coxhead and Rafiq Ahmed for assistance with Illumina RNA-sequencing, and to Javier Gilbert Jaramillo for discussions.

## Author contributions

ATH, CFH, FG, CB, JV, PL, SJC and CJAD did experiments and analysed data. SAC, WSJ, SH and CJAD obtained funding and provided supervision. ATH and CJAD conceived the study and with CFH drafted the manuscript. All authors revised the manuscript and approved the submitted version.

