## Supplementary materials for "Type I interferons govern Zika virus resistance of human macrophages and microglia"

**Extended data and supplementary material**

**Contents:**
Online Methods
Supplementary Tables S1 and S2
Supplementary Figure S1
Supplementary Figure S2
Supplementary Figure S3
Supplementary Figure S4
Supplementary Figure S5
Supplementary References

### Online Methods

#### iPSC reprogramming and culture

Human induced pluripotent stem cells (iPSC) were reprogrammed from skin dermal fibroblasts from a patient carrying a homozygous nonsense variant in *IFNAR2* (c.A311del, p.Gly104fs110X, termed *IFNAR2*$^{PT}$). Written informed parental consent was provided (Newcastle and North Tyne Research Ethics Committee Ref: 16/NE/0002). Briefly, fibroblasts were reprogrammed at passage 3-5, using Cytotune 2 Sendai virus reprogramming kit (Invitrogen) according to the manufacturer's instructions, with modifications as described[1]. QC assays on bulk banked stocks (at or near to passage 10) included flow cytometry assessment of pluripotency marker expression, RT-PCR to check clearance of cytotune vectors and Illumina SNP array to check genome integrity and identity/tracking back to the source fibroblasts (Fig S1a-c). Clones 6 and 11 were used for experiments.

The *IFNAR2*$^{WT}$ control iPSC lines SFC841-03-01[2] (WT1, Ref: STBCi044-A) and SFC856-03-04[1] (WT2, Human Pluripotent Stem Cell Registry Ref: STBCi063-A) were previously generated using the same methodology, at the James Martin Stem Cell Facility at the University of Oxford. These lines are available from the European Bank for Induced pluripotent Stem Cells (EBISC). iPSC were cultured feeder-free on Geltrex (Gibco) coated plates in mTeSR1 (Stem Cell Technologies, Canada) media, exchanged daily. Cultures were passaged using 0.5 mM EDTA (Gibco). Cryopreserved master stocks of iPSC were used and passaged no more than three times prior to differentiation to macrophages or microglia.

#### *IFNAR2* knockout (KO) using CRISPR/Cas9

The control iPSC lines SFC856-03-04 and SFC841-03-01 were used for gene editing. Two guide RNAs (gRNAs) were used to target the *IFNAR2* exon 5 splice acceptor (reference transcript: NM_001289125.3). This was predicted to cause out of frame skipping of exon 5 from all processed *IFNAR2* transcripts, leading to nonsense mediated RNA decay and absence of IFNAR2 protein expression (Fig. S4a). The guide RNA (gRNA) sequences were: CATTTTCAATAAGATGGTTG (gRNA4) and ACCGTCCTAGAAGGATTCAG (gRNA7). These gRNAs were purchased from Integrated DNA Technologies (IDT, USA) and were complexed to tracRNA, before mixing with 1.5 μg Alt-R HiFi Cas9 Nuclease (IDT, USA) to form a ribonucleoprotein (RNP) complex, according to manufacturer's instructions. The gRNA:Cas9 RNP complex was transfected into 2 x $10^5$ iPSCs in single cell suspension by electroporation (Neon transfection system, ThermoFisher Scientific) in 10 μL using the 'HiTrans' settings: 1400 v, 20 ms width, 1 pulse. iPSC were treated with 10 μL RHO kinase inhibitor Y-27632 (AbCam Biochemicals) for 3 h prior to transfection. After 3 d iPSC were single cell plated onto irradiated mouse embryonic fibroblast feeders. Single cell colonies were subsequently picked, expanded under feeder-free conditions and screened by PCR gel electrophoresis (primers available on request). One *IFNAR2*$^{-/-}$ and isogenic *IFNAR2*$^{+/+}$ pair was selected from each parental line for further validation by PCR (Fig. S4b) and capillary sequencing (not shown).

SFC856-03-04 clones were B5_2 (*IFNAR2*$^{-/-}$) and F8 (*IFNAR2*$^{+/+}$); SFC841-03-01 clones were G2 (*IFNAR2*$^{-/-}$) and G8 (*IFNAR2*$^{+/+}$). IFNAR2 expression was verified by immunoblot (Fig. S4c).

**Cells, cytokines and inhibitors**

Vero cells were cultured in complete Dulbecco's modified Eagle's medium (DMEM; Gibco) supplemented with 10% foetal calf serum (FCS; Gibco), 1% penicillin/streptomycin (Gibco) and 1% L-glutamine (Gibco). All cells were incubated in a humidified atmosphere with 5% $CO_2$ at 37°C. Cytokines/inhibitors were used at the following concentrations: human recombinant IFN-$\alpha$2b (1000 IU/ml; Intron A, Schering-Plough, USA); IFN-$\gamma$ (1000 IU/ml; Immunikin, Boehringer Ingelheim, Germany) and Ruxolitinib (10 $\mu$M; Calbiochem, USA).

**Macrophage and microglia differentiation**

iPSC were differentiated to macrophage precursor cells via embryoid bodies (EBs) using AggreWells (Stem Cell Technologies) using a published methodology with no alterations (PMID: 23951090). Media for embryoid body generation consisted of mTeSR with 50 mg/mL BMP4, 50 mg/mL VEGF and 20 mg/mL SCF. Approx. 300 EBs were split between two T175 flasks in 15 mL of medium per flask. 'Factory' medium comprised XVIVO15 (Lonza, Basel, Switzerland) with 100 ng/mL recombinant human M-CSF (Gibco), 25 ng/mL recombinant human IL-3 (Gibco), 2 nM GlutaMAX (Gibco), 50 µM 2-mercaptoethanol and 100 units/mL penicillin with 100 µg/mL streptomycin (Gibco). Precursor cells were harvested and plated into final experimental format for final differentiation to either iPS-M$\phi$ in macrophage differentiation medium (XVIVO15 with 100ng/mL recombinant human M-CSF (Gibco), 2 mM GlutaMAX (Gibco) and 100 units/mL penicillin with 100 µg/mL streptomycin) for 7 days prior to use, or iPS-microglia-like cells (iPS-MGLs) in microglia differentiation medium (Advanced DMEM/F12 + N2 Supplement (Gibco) with 100 ng/mL recombinant human IL-34 (PeproTech), 10 ng/mL recombinant human GM-CSF (Gibco), 2 mM Glutamax, 50 µM 2-mercaptoethanol and 100 units/mL penicillin with 100 µg/mL streptomycin) for 14 days, changing medium twice weekly, prior to use in experiments.

**ZIKV propagation and infection**

Two strains of ZIKV were used in this study, the Asian lineage strain H/FP/2013 (patient isolate, French Polynesia, 2013, herein ZIKV$^{FP}$) was obtained from the European Virus Archive (Ref: 001v-EVA1545) and the African lineage strain MP1751 (mosquito isolate, Uganda, 1962, herein ZIKV$^{MP}$) was obtained from the UK National Collection of Pathogenic Viruses (Ref: 1308258v). Viral stocks were propagated at low MOI in Vero cells (obtained from Professor R Randall, St Andrew's University), and viral titre determined by serial dilution and standard plaque assay. Stocks were aliquoted, clarified by centrifugation and clarified supernatants were aliquoted and frozen at -80°C, and thawed for single use. The same stocks of each virus were used for all experiments.

For experimental infection, MOI was approximated as the ratio of PFU to cells. Cells were exposed to a known titre of virus in a standard volume (50 μL for 96 well plates, 250 μL for 24 well plates, 1000 μL for 6 well plates) in macrophage or microglia medium. In parallel, cells were mock infected with medium in the same volume but without virus. At 2 hours post infection, the inoculum was removed and replaced with fresh medium (macrophage or microglia), with or without treatments (IFNs, RUX etc) as indicated, until the time required for the experiment.

**Plaque assay**

Supernatant from infected cell cultures were harvested and stored at the indicated timepoints. Aliquots of supernatant were thawed and serially diluted in DMEM with 1% FCS (Gibco), and 250 μL added to 24-well plates of confluent Vero cells. After 2 hours of incubation, 1.5% methylcellulose in DMEM with 1% FCS (Gibco) was gently added. At 72 hours post infection, media was aspirated and cells fixed with 4% formaldehyde, before being stained with 0.25% crystal violet stain, washed and plaques counted on a lightbox to determine plaque-forming units per mL (PFU/mL).

**Quantitative RT-PCR**

iPS-Mϕ were lysed in BL buffer at 24h.p.i. $ZIKV^{FP}$ MOI=10, and total RNA was extracted using the ReliaPrepTM RNA Cell Miniprep System (Promega, USA) and treated with DNase I according to the manufacturer's instructions. Quantity of purified RNA was measured spectrophotometrically (A260/A280) using a NanoDrop $One^{C}$ Microvolume UV-Vis Spectrophotometer (Thermo Fisher, MA, USA). RT-PCR was performed using the SensiFAST™ Probe No-ROX One-Step Kit (Bioline, USA) with AriaMx Real-time PCR System (Agilent Technologies, CA, USA) according to manufacturer's instructions. The following TaqMan gene expression assay (Thermo Fisher) was used: *IFNA1* (Hs03044218_g1). The primers were designed using the Roche Universal ProbeLibrary Assay Design tool (Roche, Basel, Switzerland) with the indicated UPL probes. Further details, including additional primer/probe information, are summarised in Table S1. Target gene expression was normalised to the housekeeper *GAPDH*. Each sample was run in technical duplicate. Cycling conditions were as follows: reverse transcription at 50 °C during 15 min, followed by initial polymerase activation at 95 °C for 10 min, and then 40 cycles of denaturation at 95 °C for 15 sec and annealing/extension at 60 °C for 1 min.

**RNA Sequencing**

RNA was extracted from iPS-Mϕ at 24h.p.i. $ZIKV^{FP}$ MOI=10 using BL buffer and ReliaPrepTM RNA Cell Miniprep System (Promega) according to the manufacturer's instructions. Sequencing was performed over four lanes of a single flow cell on an Illumina NextSeq (Illumina, USA) over 75 cycles. Sequencing was single ended. Raw FASTQ files were first inspected for quality using FastQC[3] and MultiQC[4]. All FASTQ files were of a very high quality, so no filtering or trimming was performed. Pseudo-alignment and transcript counting was performed and generated for the four FASTQ files for each sample using Salmon[5]. These counts are then imported into R for subsequent analysis. Transcripts were agglomerated into gene counts. Individual sample counts were merged into a single count table of samples as columns and genes as rows, with cells representing individual raw counts. Library normalisation, dispersion estimation, and log transformation was performed on the count table using the DESeq2 pipeline[6], set as default. RNA-seq data were uploaded to the Gene Expression Online (GEO) repository (accession no. pending).

**Flow Cytometry**

Macrophages were lifted using EDTA and washed with FACS buffer (PBS plus 2% FCS and 0.05% sodium azide). Cells were stained at room temperature for one hour with the following antibodies: CD45 (clone 2D1, APC-H7), CD14 (clone M5E2, BUV737), CD11c (clone B-ly6, BV421), CD86 (clone 2331, FITC, all from BD), HLA-DR (clone L243, BV650), CD11b (clone ICRF4, BV785, all from Biolegend) or the respective isotype controls. Cell viability was assessed using 7-AAD (Biolegend). After the final washing step samples were acquired on a Symphony A5 flow cytometer (BD) and results were analysed using FlowJo (Ashland, OR, USA).

**Phagocytosis Assay**

iPS-macrophages were lifted using EDTA and resuspended in fresh macrophage medium (XVIVO15 containing 1% Penicillin/Streptomycin, 1% GlutaMAX, 100 ng/mL MCSF). pHrodo red *Zymosan A* bioparticles (Invitrogen) were diluted at a concentration of 100,000 beads per ul and mixed with the iPS-macrophages at a ratio of 10:1 beads to cells. Cells were either placed on ice immediately (negative control) or incubated for 2 hours at 37$^{o}$C in a rocking incubator. Afterwards cells were immediately placed on ice to stop phagocytosis and 4',6-diamidin-2-phenylindol (DAPI; ThermoFisher) was added to assess viability. Samples were acquired without further washing steps on a Symphony A5 flow cytometer (BD Biosciences) and results were analyzed using FlowJo (Ashland, OR, USA).

**Immunoblotting**

Proteins from cell lysates were separated by 10% sodium dodecyl sulphate–polyacrylamide gel electrophoresis (SDS-PAGE) gel electrophoresis using MOPS running buffer (Thermo Fisher), and transferred to a nitrocellulose membrane (Millipore, USA) using NuPage Tris-Glycine Transfer Buffer (Thermo Fisher) for immunoblotting (antibodies see Table S2). Blots were developed with Pierce ECL Western blotting substrate (Thermo Fisher) and imaged on a LI-COR Odyssey Fc (LI-COR, NE, USA). Densitometry analysis was undertaken using ImageStudio software (version 5.2.5, Li-COR).

**Immunofluorescence**

Cells were grown on eight-well chamber slides (Millipore). Following treatment and/or infection, cells were fixed with 4% paraformaldehyde in PBS for 20 minutes at room temperature before blocking/permeabilisation with 0.5% Triton X-100 (Sigma-Aldrich, USA)/10% goat serum (Abcam, UK) in PBS for 1 hour at room temperature. Cells were incubated with mouse anti-ZIKA ENV and rabbit anti-IFITM3 specific primary antibodies (see Table S2) overnight at 4°C, then washed three times in PBS and incubated with goat anti-mouse IgG Alexa fluor 488 or goat anti-rabbit IgG Alexa fluor 555 secondary antibody (both 1 μg/ml; Thermo Fisher) for 1 hour at RT. Nuclei were stained with 4′,6-diamidino-2-phenylindole (DAPI; 0.2 μg/ml; Sigma-Aldrich). All fluorescent images were taken using an EVOS FL fluorescence microscope (Thermo Fisher) at 10X magnification using DAPI, GFP, RFP and bright-field filters. Image analysis was performed using CellProfiler software version 3.1.8 (Broad institute, USA)[7].

Images were converted to greyscale, and within DAPI images each object (nucleus) was identified using the IdentifyPrimaryObject module. In order to quantify proteins within the whole cell including the cytoplasm, cells were identified within bright-field images by propagating out from each nucleus using the IdentifySecondaryObjects module, to create a mask for each cell. Predefined thresholds for GFP (Env protein) and RFP (IFITM3) were used to classify Zika positive and/or ISG productive cells, and the average value of at least n=4 images per well was used for analysis. A total of 58 sets of images (since all biological replicates were made up of at least n=3 technical replicate wells) with a median (IQR) of 255.5 (162 – 513.25) cells per image processed.

**Cell viability Assay**

Cells in 96 well plates were infected at the indicated MOI for 72 hour, or pre-treated as indicated for 16 hours before infection and imaging. Live cell imaging solution containing 3 drops/mL of ReadyProbes Cell Viability Blue/Red Imaging Kit (Invitrogen) was added 30 minutes before imaging to assess cell viability. Images were obtained using an EVOS FL fluorescence microscope (Thermo Fisher) using DAPI and RFP filters at 10x magnification. Image analysis was performed using CellProfiler version 3.1.8 (Broad institute, USA). DAPI and RFP images were converted to greyscale and objects identified using IdentifyPrimaryObject modules. DAPI objects (corresponding to Hoechst nuclear staining) overlapping with RFP objects (corresponding to propidium iodide staining) were classified as dead cells, while non-overlapping DAPI objects were counted as live cells. All conditions were performed in technical duplicate and the average value of n=6 images per well was used for analysis. All experiments were performed at least three times. The cell profiler pipelines used are available at https://github.com/aidanhanrath/Cellprofiler.

**Statistical Analysis**

Statistical analysis was performed using GraphPad Prism 9 software (GraphPad Software, CA, USA). Values are presented as mean ±SD of at least three independent experimental replicates. Statistical significances between two groups of data were determined using an unpaired, two-tailed Student's t-test. Statistical analysis of data sets with two independent variables was carried out using a two-tailed one way analysis of variance (ANOVA) with a post-hoc Sidak's test to account for multiple comparisons. $P<0.05$ was considered statistically significant.

| Gene | UPL probe | Forward sequence | Reverse sequence |
|---|---|---|---|
| *IFNB* | #25 | CGACACTGTTCGTGTTGTCA | GAAGCACAACAGGAGAGCAA |
| *IFNL1* | #75 | GGGACCTGAGGCTTCTCC | CCAGGACCTTCAGCGTCA |
| *GAPDH* | #87 | TGGTATCGTGGAAGGACTCA | GCCATCACGCCACAGTTT |

**Supplementary Table S1. QRT-PCR primer/probe sequences.** Note *IFNA1* was assessed using a commercial kit (Hs03044218_g1, Thermofisher).

| Antibody | Host | Working dilution | Source | Code |
|---|---|---|---|---|
| IFNAR2 | Sheep | 1:200 | R&D systems | AF7014 |
| IFITM3 | Rabbit | 1:10000 | ProteinTech | 11714-1-AP |
| RSAD2 | Rabbit | 1:1000 | CST | 13996 |
| ISG15 | Rabbit | 1:1000 | CST | 2743 |
| STAT2 | Mouse | 1:2000 | SCB | sc-1668 |
| pSTAT2 | Rabbit | 1:2000 | CST | 8841 |
| STAT1 | Rabbit | 1:1000 | CST | 9172 |
| pSTAT1 | Rabbit | 1:1000 | CST | 7649 |
| Tmem119 | Rabbit | 1:100 | Abcam | Ab185333 |
| Iba1 | Goat | 1:500 | Abcam | Ab5076 |
| Zika Envelope | Mouse | 1:5000 | BioFront Tech | BF-1176-56 |
| α-tubulin | Mouse | 1:10,000 | CST | 3873 |
| GAPDH | Rabbit | 1:10,000 | CST | 5174 |
| Anti-rabbit HRP-conjugated | Goat | Various | CST | 7074 |
| Anti-mouse HRP-conjugated | Horse | Various | CST | 7076 |

**Supplementary Table S2. Immunoblotting and immunofluorescence antibodies.** CST = Cell Signalling Technologies; HRP = horseradish peroxidase; SCB = Santa Cruz Biotechnology.

### Supplementary Figures

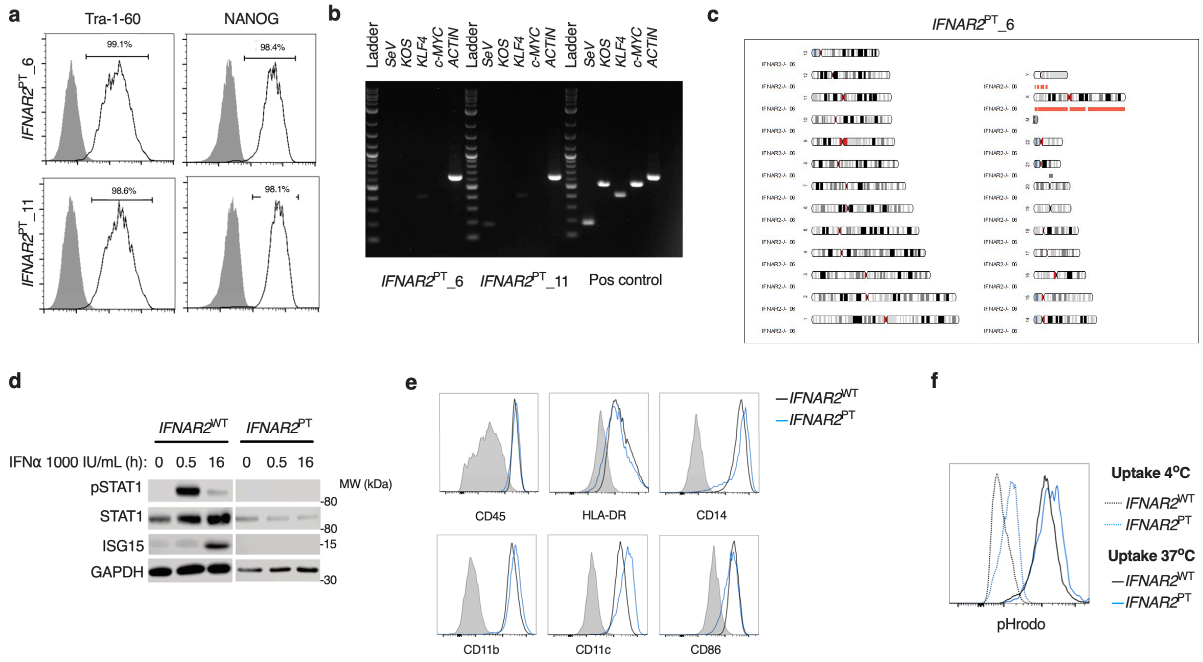


**Figure S1. A model of IFNAR2-deficient human iPS-macrophages (*IFNAR2*$^{PT}$ iPS-Mϕ).**

(a) Expression of pluripotency markers by *IFNAR2*$^{PT}$ iPSC clones 6 and 11 (flow cytometry).

(b) PCR showing clearance of cytotune Sendai vector from *IFNAR2*$^{PT}$ iPSC clones 6 and 11.

(c) Karyogram produced from SNP array showing no gross abnormalities in the previously unpublished *IFNAR2*$^{PT}$ iPSC clone 6. Red bars indicate loss or single copy, grey indicates loss of heterozygosity on chromosome 21 in the region of *IFNAR2* (representative of data in *IFNAR2*$^{PT}$ clone 11).

(d) Immunoblot of IFN-I signalling in IFNα2b (1000 IU/mL) treated *IFNAR2*$^{PT}$ (clone 11) and *IFNAR2*$^{WT}$ (WT1) iPS-Mϕ, representative of n=3 repeat experiments in *IFNAR2*$^{PT}$ clone 11 and WT2.

(e) Expression of macrophage surface markers in *IFNAR2*$^{PT}$ (clone 6) and *IFNAR2*$^{WT}$ (WT1) iPS-Mϕ by flow cytometry, representative of repeat experiments in clone 11 and WT2.

(f) Phagocytic uptake of Zymosan pHrodo particles in *IFNAR2*$^{PT}$ (clone 6) and *IFNAR2*$^{WT}$ (WT1) iPS-Mϕ, representative of repeat experiments in clone 11 and WT2.

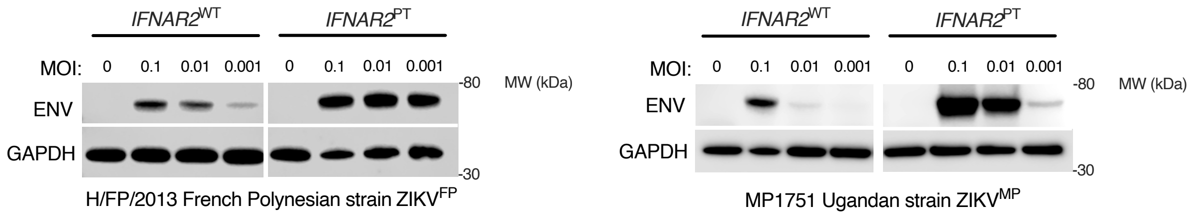


**Figure S2. Enhanced ZIKV$^{HFP}$ and ZIKV$^{MP}$ replication in *IFNAR2*$^{PT}$ iPS-Mϕ.**
Immunoblot of ENV and GAPDH expression in *IFNAR2*$^{PT}$ and *IFNAR2*$^{WT}$ iPS-Mϕ, 72h.p.i. post infection, representative of n=3 repeats in *IFNAR2*$^{PT}$ clone 11 and *IFNAR2*$^{WT}$ WT2.

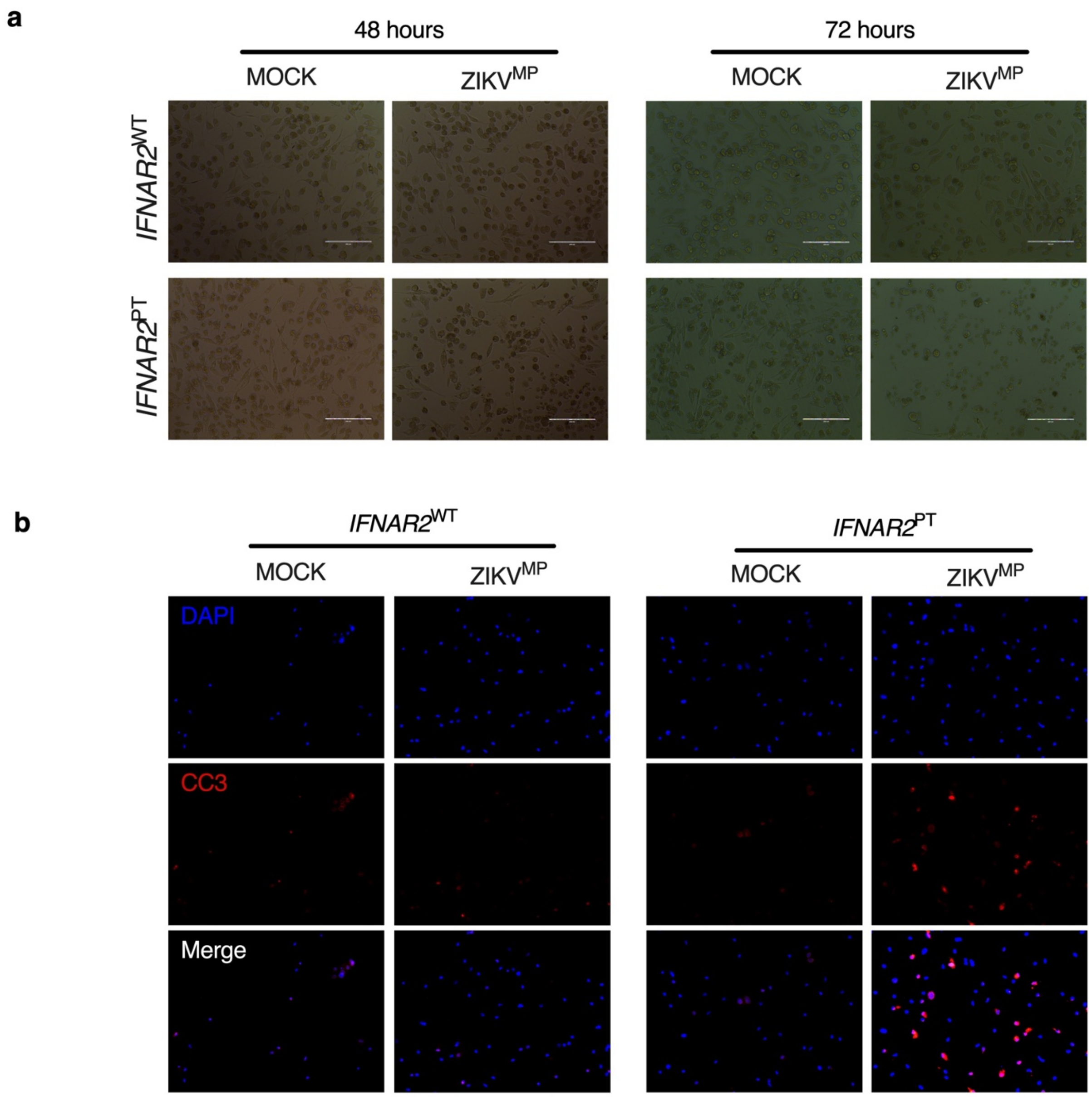


**Figure S3. Morphological changes of ZIKV-induced cell death in *IFNAR2*$^{PT}$ iPS-M$\phi$.**

(a) Progressive cytopathicity in *IFNAR2*$^{PT}$ (clone 6) but not *IFNAR2*$^{WT}$ (WT2) iPS-M$\phi$, with features of cell shrinkage and membrane blebbing, representative of n=3 repeat experiments. Error bars = 200 $\mu$m.

(b) Immunofluorescence analysis of cleaved caspase 3 (CC3) at 48h.p.i., representative of n=2 repeat experiments.

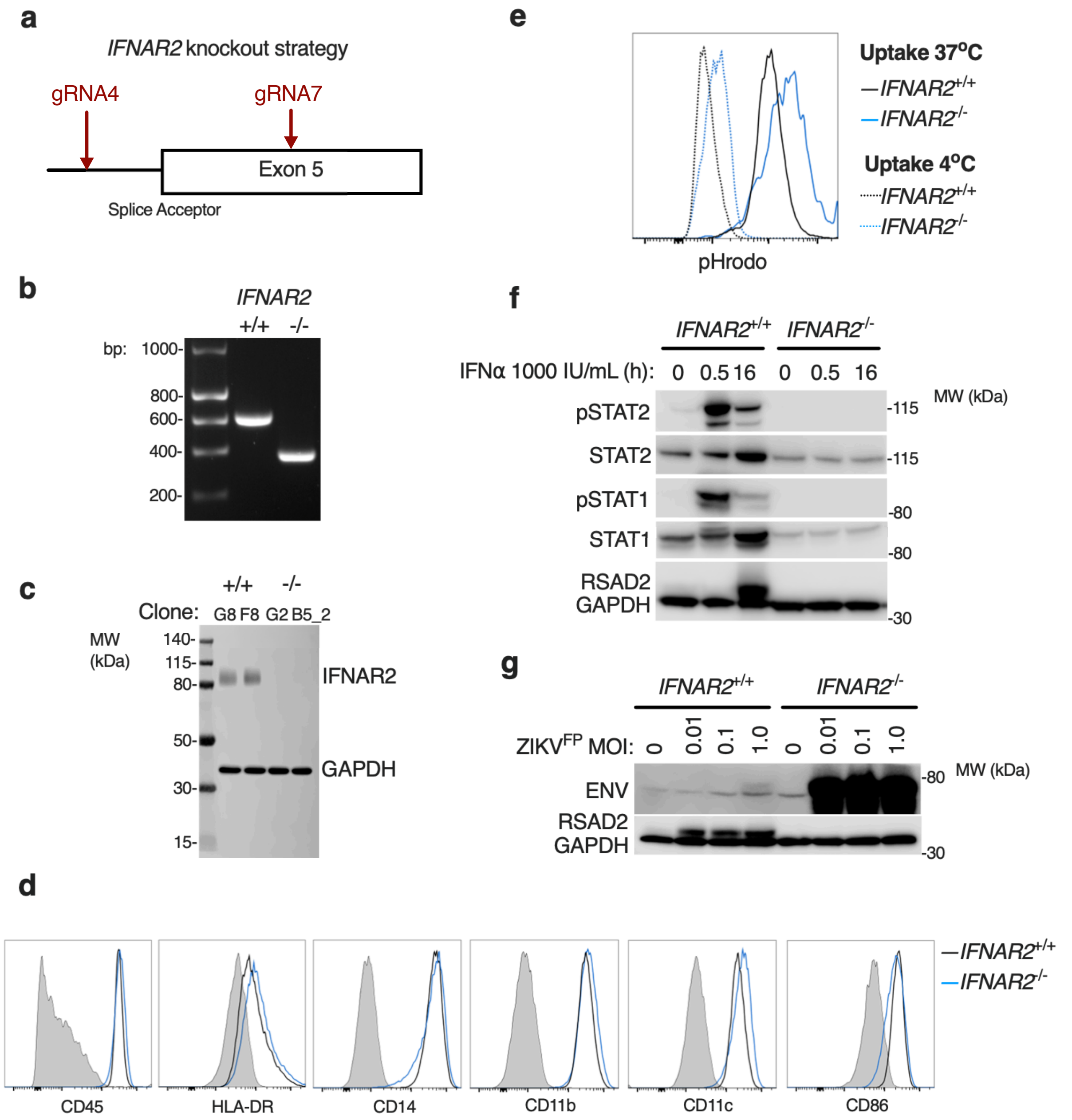


**Figure S4. CRISPR/Cas9 knockout of *IFNAR2.***

(a) CRISPR guide design.

(b) PCR demonstrating *IFNAR2* exon 5 splice acceptor site excision in iPSC clones (G8 and G2).

(c) Immunoblot of IFNAR2 and GAPDH expression, demonstrating IFNAR2 ablation in iPS-Mϕ clones (G8, F8, G2, B5_2). Representative of n=2 repeat experiments.

(d) Expression of macrophage surface markers in *IFNAR2*$^{-/-}$ (B5_2) and *IFNAR2*$^{+/+}$ (F8) iPS-Mϕ by flow cytometry, representative of n=2 repeat experiments in F8/B5_2.

(e) Phagocytic uptake of Zymosan pHrodo particles in *IFNAR2*$^{-/-}$ (B5_2) and *IFNAR2*$^{+/+}$ (F8) iPS-Mϕ by flow cytometry, representative of n=2 repeat experiments in F8/B5_2.

(f) Immunoblot of IFN-I signalling in IFN$\alpha$2b (1000 IU/mL) treated *IFNAR2*$^{-/-}$ and *IFNAR2*$^{+/+}$ iPS-M$\phi$, representative of n=3 repeats in F8/B5_2.

(g) Immunoblot of ENV, ISG15 and GAPDH expression in *IFNAR2*$^{-/-}$ and *IFNAR2*$^{+/+}$ iPS-M$\phi$, 72h.p.i. post infection, representative of n=3 repeats in F8/B5_2.

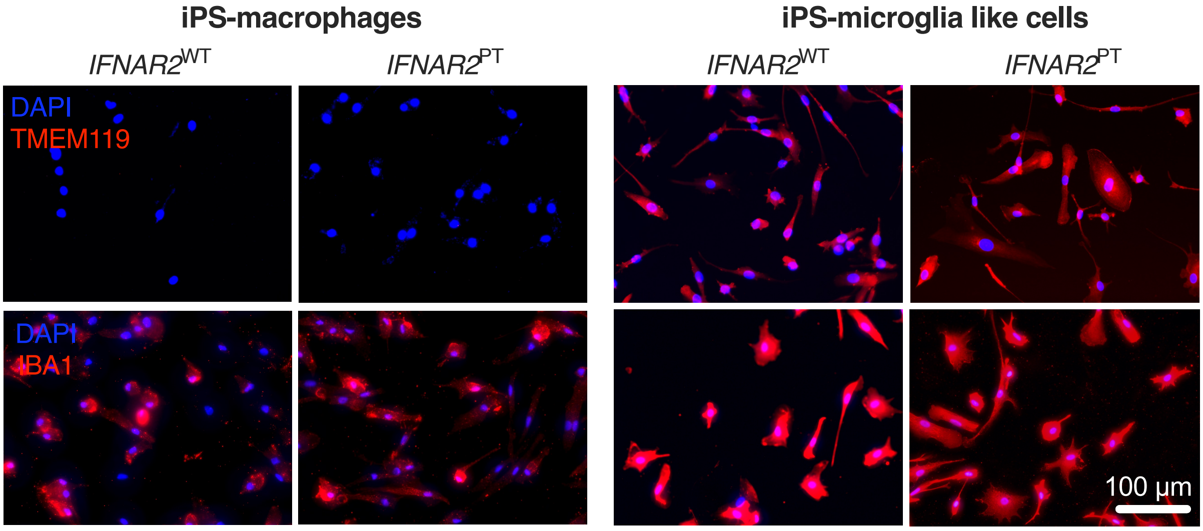


**Figure S5.** Microglial marker expression in iPS-Mϕ and iPS-microglia-like cells (iPS-MGLs). Immunofluorescence analysis of microglial markers IBA1 and TMEM119 in *IFNAR2*$^{PT}$ (PT clone 11) and *IFNAR2*$^{WT}$ (WT2) iPS-Mϕ and iPS-MGLs, representative of n=3 repeat experiments in G2, B5_2, G8 and F8 lines.
